# Ubp3 mediates dynamic deubiquitination of mitochondria upon induction of mitophagy

**DOI:** 10.64898/2026.09.17.752381

**Authors:** Kai Mayor Völtzke, Lars Bostelmann-Arp, Christina Behrendt, Mareike Breuer, Simon Vesper, Katja Bendrin, Lenka Geerkens, Melissa Vazquez-Carrada, Lasse van Wijlick, Ursula Fleig, Michael Feldbrügge, Lutz Schmitt, Andreas S. Reichert

**Author notes:** To whom correspondence should be addressed: Andreas S. Reichert.

## Abstract

Ensuring quality and maintenance of mitochondria within eukaryotic cells is important for cellular fitness. One critical pathway that needs to be tightly controlled to achieve this is mitophagy, which is negatively regulated by the deubiquitinase Ubp3 in *Saccharomyces cerevisiae.* Here we show that beyond this established role of Ubp3, it is critical to mediate substantial temporal changes of the mitochondrial ubiquitin landscape during mitophagy. Isolated mitochondria displayed extensive ubiquitination under steady-state conditions, whereas rapamycin-mediated induction of mitophagy/autophagy led to a progressive elimination of mitochondrial ubiquitination. A systematic screening analysis revealed that Ubp3 and Ubp4 are crucial regulators of rapamycin-induced mitochondrial deubiquitination. Deletion of Bre5, a cofactor of Ubp3, or of the *UBI4* gene, encoding Ubi4 required for mitophagy, did not impair deubiquitination of mitochondria during mitophagy, indicating that Ubp3 has additional functions. We further showed that Ubi4 acts epistatic to Ubp3 for regulating mitophagy, supporting that mitophagy does not simply depend on the extent of mitochondrial ubiquitination, but rather on specific ubiquitinated substrates. Consistent with the role of Ubp3 *in vivo*, purified Ubp3 or its catalytic domain alone were sufficient to efficiently deubiquitinate isolated mitochondria *in vitro* in a Bre5-independent manner. Together, these findings highlight Ubp3 as a prominent rheostat of mitochondrial ubiquitin remodelling, revealing mitochondrial ubiquitin storage and its dynamic release as another layer of stress regulation by ubiquitin-dependent quality control pathways.

## Introduction

Mitochondria are essential organelles in eukaryotic cells that cannot be generated *de novo*. Consequently, their continuous biogenesis and division of pre-existing organelles, and their quality control and degradation represent a network that needs to be tightly regulated (Kondadi and Reichert, 2024). Conversely, removal of damaged organelles is as vital as preserving a healthy subset for propagation. Cells have evolved elaborated mechanisms for the orchestrated removal of organelles via proteasomal degradation and autophagy. The selective removal of mitochondria via autophagy is termed mitophagy and occurs either via receptor-mediated mitophagy or by ubiquitin-dependent mitophagy (Abeliovich, 2023; Zimmermann and Reichert, 2017). In *Saccharomyces cerevisiae* the only known mitophagy-related receptor protein Atg32 has been studied intensively. Atg32 is activated and accumulates on the outer membrane of mitochondria where it recruits via interaction with Atg11 and Atg8 the autophagosome precursor thus initiating organelle elimination (Abeliovich, 2023). In humans, mitophagy is known to prevent accumulation of impaired mitochondria via ubiquitin-mediated mitophagy requiring the PTEN-induced putative kinase 1 (PINK1)- and E3-ubiquitin ligase PARKIN-mediated ubiquitination, two factors associated with the development of neurodegenerative diseases such as Parkinson’s disease (Narendra et al., 2010; Vives-Bauza et al., 2010; Youle and Narendra, 2011). To date no PINK1 or PARKIN homologs have been identified in yeast but one study showed that transforming yeast cells with human PARKIN results in increased mitochondrial degradation (Pereira et al., 2015). Moreover, in a genome-wide screen for regulators of mitophagy the subunits of the Ubp3/Bre5 deubiquitinase complex were identified as negative regulators of mitophagy (Müller et al., 2015). A recent study revealed that the polyubiquitin-encoding gene *UBI4*, or more precisely, a C-terminally extended form of ubiquitin (CxUb/Ub77N), is crucial for mitophagy in budding yeast (Altin et al., 2025). Together these studies clearly show that the machinery for ubiquitin-mediated regulation of mitophagy exists also in budding yeast and may share conserved mechanisms with mammalian mitophagy.

In *S. cerevisiae*, the ubiquitinome acts as a central signalling hub that gets dynamically altered upon stress, orchestrating multiple cellular quality control pathways (Dikic, 2017; Sheng et al., 2024). Treatment with the starvation mimetic rapamycin, an inhibitor of the mechanistic target of rapamycin (mTor), rapidly alters the ubiquitination state of yeast proteins (Iesmantavicius et al., 2014; Kanki et al., 2011). The dynamic effect of rapamycin treatment on mitochondrial ubiquitination was unknown. Ubiquitin is synthesized either as a ribosomal fusion protein encoded by *UBI1/UBI2/UBI3* or as a polyubiquitin precursor by *UBI4*, which are subsequently matured by deubiquitinases (DUBs). The ubiquitin code (Komander and Rape, 2012) is a highly complex system and gets regulated by a plethora of so termed *writers* (E1 activating enzyme, E2 conjugating enzymes, and E3 and E4 ligases) and *erasers* or deubiquitinating proteases (DUBs). DUBS are divided into ubiquitin-specific proteases (Ubps) and ubiquitin carboxy-terminal hydrolases (UCHs). The localization and specific targets for different DUBs have been subject to investigation but the role of DUBs in mitochondrial quality control is not well understood. Ubp2 and Ubp12 have been implicated in mitochondrial dynamics, and antagonistically promote fusion and fission, respectively (Anton et al., 2013; Biswas and D’Silva, 2025). Ubp8 has been shown to promote mitochondrial respiration, however, this regulation occurs via promoting gene transcription and thus is unlikely to happen by directly deubiquitination of mitochondrial proteins (Leo et al., 2018). Kinner and Kölling showed that Ubp16 is mitochondria membrane-anchored, yet apparently does not play a role in mitochondrial quality control (Kinner and Kölling, 2003). Finally, Ubp3 together with its proposed positive regulator Bre5 was shown by us to dynamically relocate from the cytoplasm to mitochondria upon treatment with rapamycin and was well established to inhibit mitophagy (Müller et al., 2015). Consequently, it has been hypothesized that the underlying mechanism of regulating mitophagy by Ubp3 is deubiquitination of mitochondria, however, this was not experimentally demonstrated (Behrendt and Reichert, 2016; Müller et al., 2015). It was also not known to which extent mitochondria are ubiquitinated under non-stressed conditions and whether the mitochondrial ubiquitin profile changes over time. If so, whether a specific DUB or a network of different DUBs catalyse the remodelling remains to be elucidated.

Here, we studied the ubiquitination status of isolated mitochondria during a time course of rapamycin treatment to characterize the dynamic remodelling of the ubiquitin profile of mitochondria under starvation-like conditions. To identify potential regulators of this dynamic process, we investigated the role of the major DUBs existing in budding yeast on the deubiquitination of mitochondrial proteins. Specifically, in this study we delineated the function of Ubp3 in mitochondrial ubiquitin dynamics and propose Ubp3 as a major factor in mitochondrial ubiquitin remodelling.

## Results

## Ubiquitin dynamically translocates from mitochondria to the cytosol upon induction of autophagy with rapamycin

We have recently shown that mitophagy depends on the polyubiquitin-encoding gene *UBI4* in *S. cerevisiae* (Altin et al., 2025). To address where this protein is specifically located before and during mitophagy, we expressed UBI4-GFP, a C-terminally GFP-tagged variant of UBI4, in *S. cerevisiae* and determined its subcellular location upon addition of rapamycin, an established inducer of mitophagy. In the absence of rapamycin, most cells showed that UBI4-GFP is located both at mitochondria and the cytosol, whereas few cells showed a cytosolic-only signal. Upon addition of rapamycin the proportion of cells showing a mitochondrial signal is rapidly reduced within 24 hours while correspondingly the cytosolic signal increases (Fig 1AB). To validate this, western blot analysis before and after subcellular fractionations was performed. After cells were either treated or not with rapamycin for 24 hours we separated total cellular lysates into cytosolic and mitochondrial fractions. We observed that the corresponding full-length UBI4-GFP (∼70 kDa) band was exclusively present in the mitochondrial fraction, and that its level was moderately reduced after rapamycin treatment (Fig. 1C). Conversely, truncated forms of UBI4-GFP as well as released GFP (∼27 kDa) were mostly found in the cytosolic fractions, which were apparently increased when cells have been treated with rapamycin. Overall, these experiments suggest that the polyubiquitin encoded by UBI4 is located at mitochondria and is released to the cytosol, in particular upon induction of mitophagy. In this context, it is interesting to note that apparently only full-length UBI4-GFP is initially stably bound to mitochondria and processing of this polyubiquitin precursor appears to occur rapidly, even in the absence of rapamycin, yet is enhanced after addition of rapamycin.

**Figure 1:**
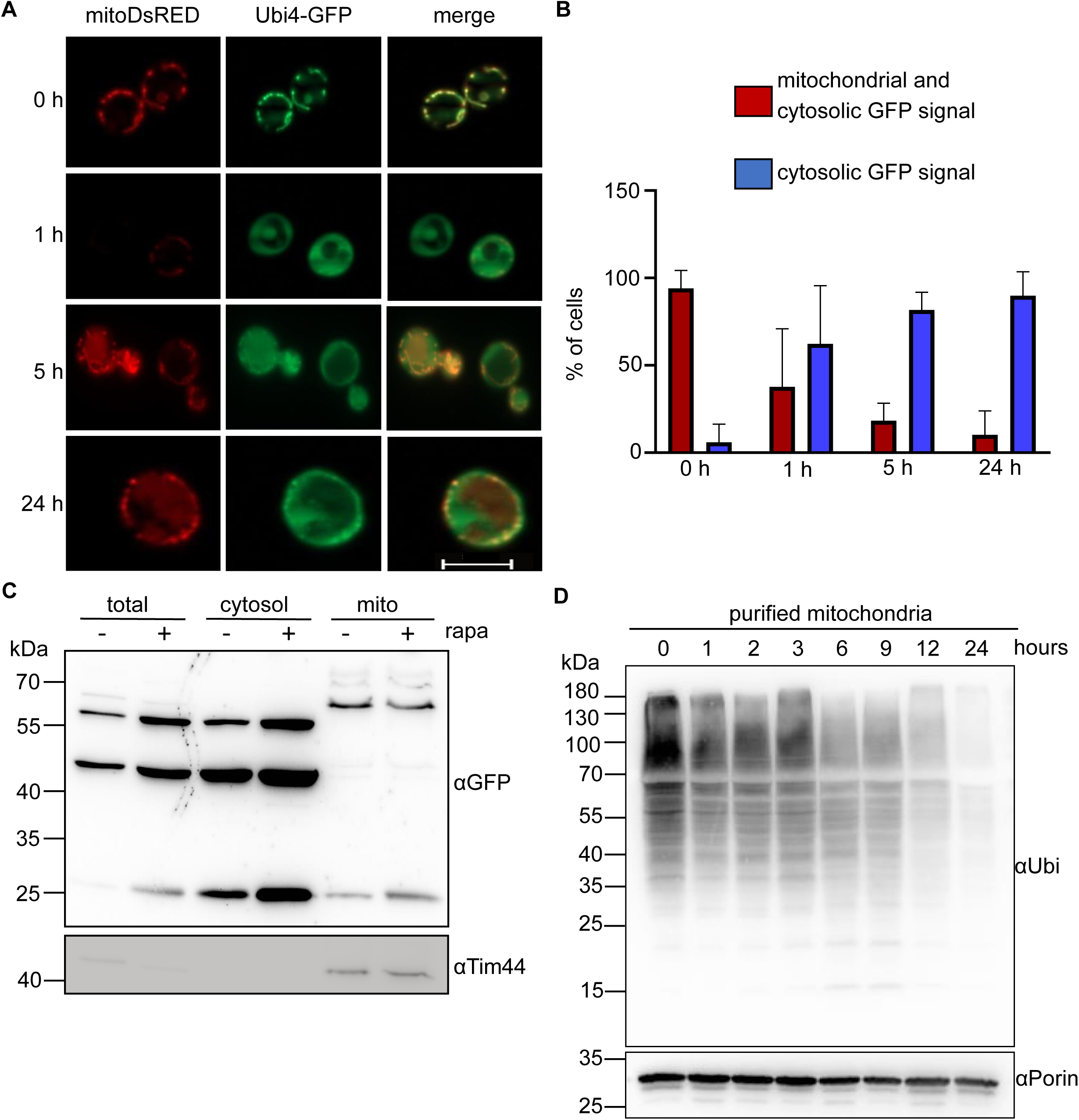
Ubiquitination profile of mitochondria is dynamically remodelled upon induction of mitophagy. (A and B) Strains expressing mitoDsRED and C-terminally GFP tagged Ubi4 were analysed by fluorescence microscopy. (A) Representative images of cells treated with rapamycin for 1, 5 or 24 hours, or left untreated as control are shown. (B) Cells from three independent experiments were analysed and the percentage of cells with Ubi4-GFP localization with a mitochondrial or cytosolic and mitochondrial GFP signal were quantified. (C) Fractionation of the same strain treated 24 hours with 1 µM rapamycin or left untreated. Total lysate, cytosolic fraction and purified mitochondria were analysed by SDS PAGE and western blotting using an anti-GFP antibody to compare the different levels and length of Ubiquitin-GFP fusion proteins between treated and untreated cells and the different cellular localizations. Mitochondrial protein Tim44 served as a loading control. (D) Purified mitochondria from WT cells were incubated with rapamycin for a time course of 0.5, 1, 2, 3, 6 or 24 hours, or left untreated. The ubiquitination status of the mitochondria was analysed by SDS-PAGE and western blotting using an anti-ubiquitin antibody. Mitochondrial protein Porin served as a loading control.

### Mitochondria harbour a significant pool of ubiquitin that is rapidly lost upon induction of autophagy

Intrigued by these observations, we next asked whether mitochondria in general harbour a substantial pool of ubiquitin and whether this, if present, is dynamically modulated upon induction of autophagy or mitophagy. For this, we isolated mitochondria from non-treated and from cells treated with rapamycin over a time course up to 24 hours (Fig. 1D). Indeed, we observed that non-treated, isolated mitochondria show a broad and extensive signal for ubiquitinated proteins suggesting that numerous mitochondrial proteins or mitochondria-associated proteins are present at the surface of this organelle under steady-state conditions. Moreover, an extensive dynamic reduction of the ubiquitination profile of mitochondria over time upon treatment with rapamycin was observed, being nearly complete after 24 hours. In sum, we conclude that mitochondria contain a substantial amount of ubiquitin which is released to the cytosol. The broad range of molecular weights observed for ubiquitinated mitochondrial proteins strongly suggests that multiple proteins harbour one or more ubiquitin moieties and that those are deubiquitinated over time upon autophagy induction.

### Deubiquitination of mitochondria is strongly dependent on the deubiqitinases Ubp3 and Ubp4 but not on Bre5 or Ubi4

To determine, which DUBs in *S. cerevisiae* are required for deubiquitinating mitochondria we performed a knock-out screen using a selection of 16 strains each lacking one known DUB. Cells from these strains were grown under control or rapamycin treated conditions and the overall ubiquitination status of purified mitochondria was analysed as previously. In strains lacking Ubp3 or Ubp4/Doa4 deubiquitination of mitochondria upon rapamycin treatment was strongly impaired compared to the wild-type control (Fig. 2AB), whereas only minor changes were observed for all other tested deletion strains. We have shown earlier that Ubp3 inhibits mitophagy and dynamically translocates to this organelle upon induction of mitophagy by rapamycin. This together with our new findings are consistent with the idea that deubiquitination of mitochondria limits excessive mitophagy and ensures maintenance of this essential organelle under stress. To follow this in more detail, we decided to focus on Ubp3 and asked whether Bre5, a cofactor of Ubp3, is needed for mitochondrial deubiquitination. Deletion of Bre5 did not prevent rapamycin-induced deubiquitination of mitochondria (Fig. 2C), which is consistent with our recent finding that Ubp3 enzyme activity does not strictly require its cofactor Bre5 *in vitro* (Bostelmann-Arp et al., 2026). The polyubiquitin-encoding gene *UBI4* is neither required for the formation of a mitochondrial ubiquitin pool under steady-state conditions nor for its dynamic deubiquitination upon addition of rapamycin (Fig. 2C), indicating that global ubiquitination of mitochondria is not sufficient to promote mitophagy and that rather specific, possible only very few, substrates need to be ubiquitinated for mitophagy in baker’s yeast. To further dissect a possible functional link between *UBP3* and *UBI4*, we asked whether the inhibitory role of Ubp3 during mitophagy is dependent on *UBI4*. For that, we quantified mitophagy using an established assay in strains lacking Ubp3, Ubi4, or both and compared it to a wild-type strain. We observed that that deletion *UBP3* does not increase mitophagy when *UBI4* is lacking as well (Fig. 3A), demonstrating that UBI4 is epistatic to Ubp3. Given that deletion of UBI4 does not prevent overall ubiquitination of mitochondria corroborates the view that *UBI4* is needed to ubiquitinate specific substrates during mitophagy rather than being needed for overall mitochondrial ubiquitination, as depicted in our model (Fig. 3D). One possible specific ubiquitinated factor, the outer membrane protein Fzo1, was recently shown to be required for mitophagy (Altin et al., 2025). Ubp3 has been shown to negatively regulate mitophagy (Müller et al., 2015). Yet, whether this is actually dependent on the catalytic activity of Ubp3 was not tested so far, prompting us to mutate the catalytic cysteine in the active site to alanine (C469A). Next to a vector control we expressed this variant or the wild-type Ubp3 in the *Δubp3* background and quantified mitophagy after rapamycin induction. Indeed, disrupting the catalytic site of Ubp3 resulted in the loss of mitophagy inhibition normally exerted by Ubp3 (Fig. 3B). Overall, we conclude that the enzymatic activity of Ubp3 as a DUB is required for inhibiting mitophagy and that Ubp3 can act independently of Bre5 in deubiquitinating mitochondria *in vivo*.

**Figure 2.**
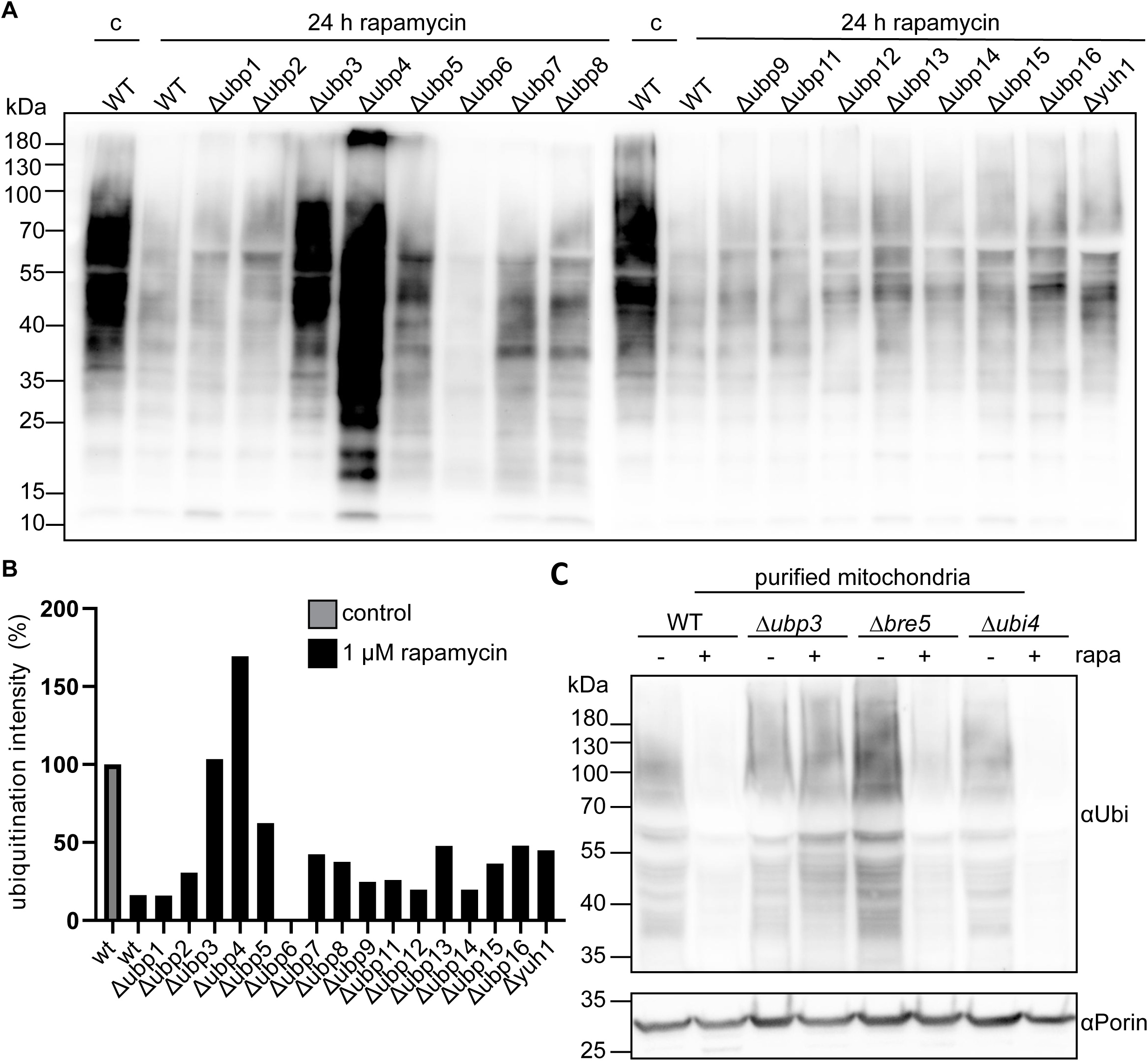
Ubp3 and Ubp4 are the prominent DUBs acting at mitochondria upon treatment with rapamycin. (A) Ubiquitin levels of mitochondria were analysed in wildtype (WT) cells or cells disrupted for UBP or UCH, as depicted, after treatment with 1 µM rapamycin for 24 hours. Deubiquitination of mitochondria was assessed by SDS–PAGE and western blotting using an anti-Ubiquitin antibody. (B) The quantification of ubiquitin intensities shown in a bar graph. (C) Ubiquitin levels of mitochondria were analysed in wildtype (WT) cells or cells disrupted for either Ubp3 *(Δubp3)*, Bre5 *(Δbre5)* or Ubi4 *(Δubi4)* after treatment with 1 µM rapamycin (black) for 24 hours or as a control without rapamycin (grey). The ubiquitination status of the mitochondria was analysed by SDS-PAGE and western blotting using an anti-Ubiquitin antibody. Mitochondrial protein Porin served as a loading control.

**Figure 3:**
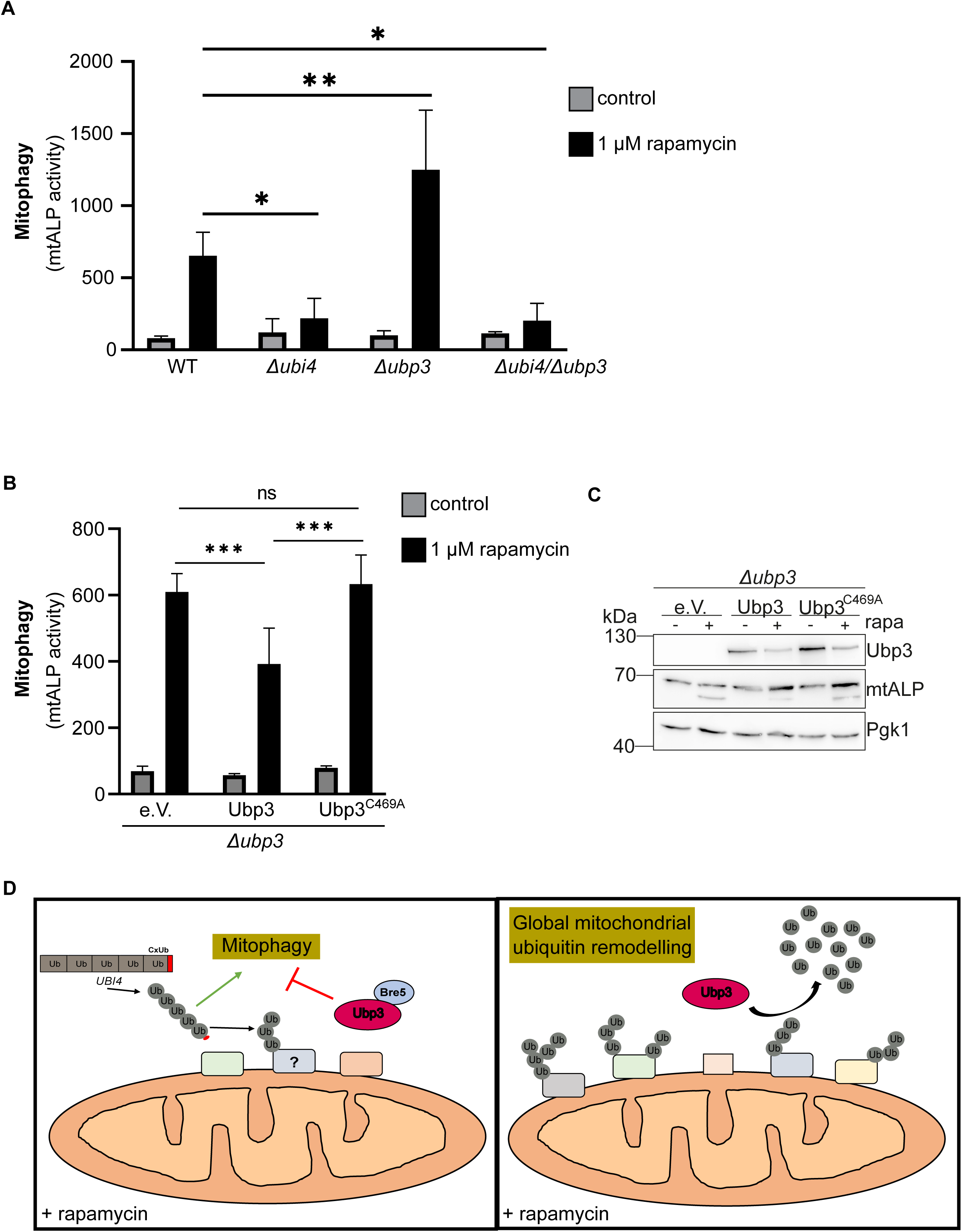
Ubp3 controls global mitochondrial ubiquitination profile whereas specific ubiquitination signals mitophagy/Ubi4 is epistatic to induction of mitophagy (?) (A) Mitophagy was assessed in wildtype (WT) cells or cells disrupted for either Ubi4 *(Δubi4)*, Ubp3 *(Δubp3)*, or Ubi4 and Ubp3 (Δubi4Δubp3). (B and C) Cells disrupted for Ubp3 (Δubp3) were transformed with either an empty vector or a vector coding for Ubp3 or a catalytic inactive Ubp3 mutant (Ubp3) and induction of mitophagy was analysed and quantified. (C) Western blot analysis of one representative experiment with indicated antibodies. (D) Schematic overview of how Ubi4 and Ubp3 regulate different aspects of mitochondrial ubiquitination and mitophagy. (A, B and C) Strains were either treated with 1 µM of rapamycin for 24 hours or left untreated and mitophagy was measured with the standard mtALP assay in strains deleted for endogenous PHO8 expressing mtALP. Mean ± SD of at least three independent experiments normalized to control is shown. Significance levels p< 0.05 (∗), p<0.01 (∗∗) or p<0.001 (∗∗∗) using a two-way ANOVA test are indicated.

### Purified Ubp3 is able to deubiquitinate isolated mitochondria in a Bre5-independent manner in vitro

These findings *in vivo* raise the question whether Ubp3 directly acts at the mitochondrial surface on mitochondrial ubiquitinated proteins or, alternatively, acts more indirectly on non-mitochondrial substrates in a pathway required for mitochondrial deubiquitination. Recently, purified Ubp3 has been shown by us to act as a deubiquitinase *in vitro* acting on three specific substrates, namely UB-AMC, Ubi-GFP and CxUb (Ub77N), which occurred independent of its proposed cofactor (Bostelmann-Arp et al., 2026). Also, the catalytic domain of Ubp3 (Ubp3_CD_) alone, lacking the binding domain for Bre5, was shown to be sufficient for efficient catalysis. To determine, whether purified full-length Ubp3 (Ubp3-FL) can directly deubiquitinate isolated mitochondria, we incubated mitochondria from *S. cerevisiae* with increasing amounts of Ubp3-FL. Indeed, the level of mitochondrial ubiquitin was reduced in a dose-dependent manner *in vitro* (Fig. 4A). Next, in addition to Ubp3-FL (FL) we used the catalytic domain of Ubp3 (CD), and as controls an empty vehicle (V) or the catalytic inactive domain of Ubp3 (CD C469A). We observed that the catalytic domain of Ubp3 is sufficient for deubiquitination of isolated mitochondria and occurred to a similar extent as the full-length Ubp3 protein (Fig. 4BC). The catalytic inactive mutant, as expected, was not able to deubiquitinate mitochondria. Moreover, adding purified Bre5 to Ubp3-FL (FL) or to the catalytic domain of Ubp3 (CD) did not enhance mitochondrial deubiquitination *in vitro* (Fig. 4D), consistent with the earlier notion that Ubp3 activity is not dependent on Bre5. We further tested whether mitochondria from other organisms are also ubiquitinated under steady-state conditions and could act as a substrate for purified Ubp3-FL or Ubp3-CD. We isolated mitochondria from human HEK cells, *S. pombe* or *U. maydis* and performed the same type of experiment. This revealed that also mitochondria from these organisms show an extensive level of ubiquitinated proteins which can be reduced by Ubp3-FL and Ubp3-CD, but not by Ubp3-CD C469A (SFig. 1). Overall, we conclude that mitochondrial ubiquitination is conserved across a broad range of species and that mitochondria can be directly deubiquitinated by purified Ubp3 from *S. cerevisiae* in a Bre5-independent manner.

**Figure 4:**
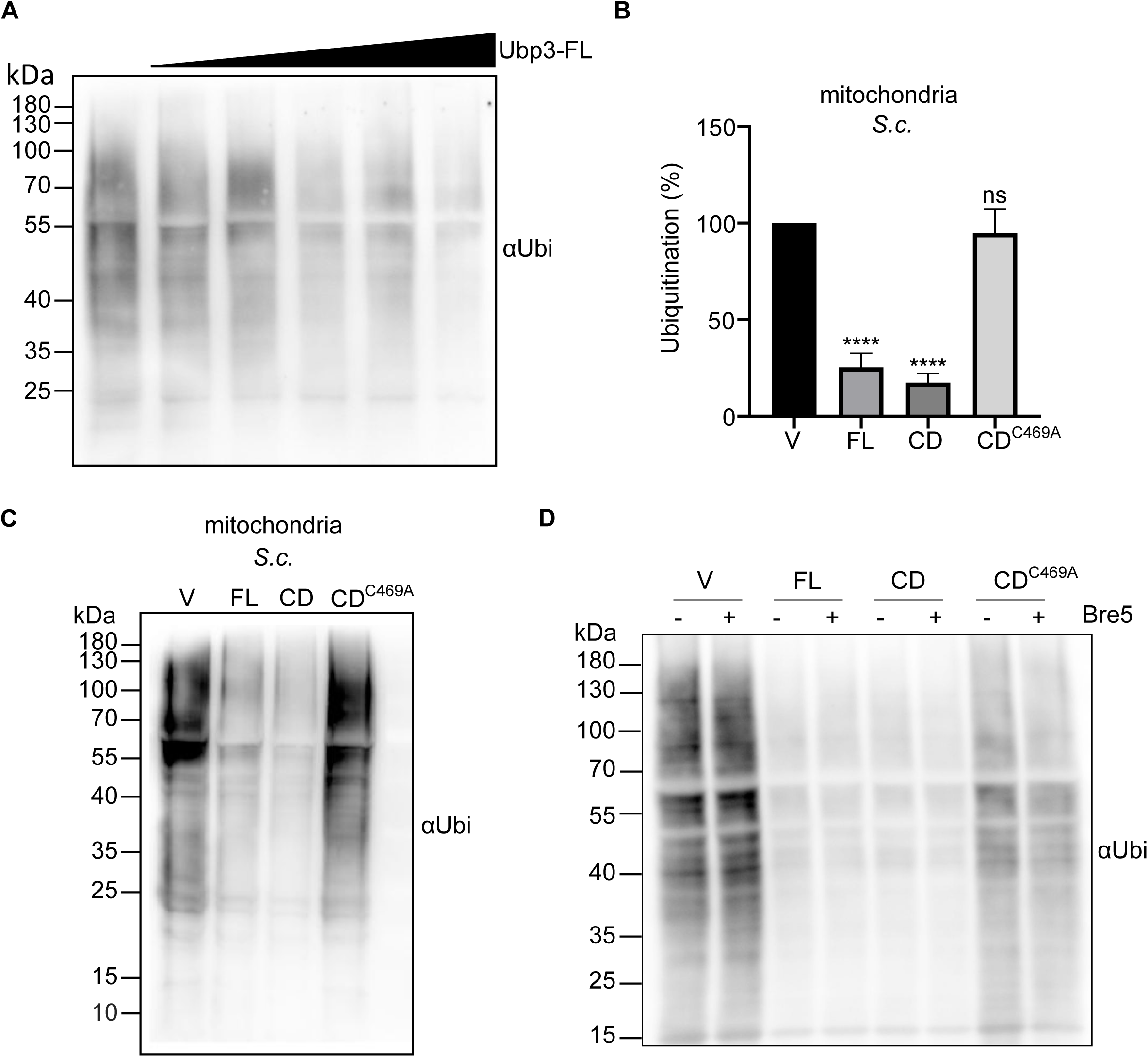
Isolated Ubp3 deubiquitinates purified mitochondria ex vivo independently of Bre5. (A) Purified mitochondria from wild-type *S. cerevisiae* were incubated with increasing doses of purified full length Ubp3. (B and C) Mitochondria isolated from wild-type *S. cerevisiae* were incubated with either vehicle control (V), or purified proteins full length Ubp3 (FL), the catalytic domain of Ubp3 alone (CD), the catalytic domain of Ubp3 carrying the inactivating C469A mutation (CD^C469A^). A quantification of the blots (B) and a representative experiment (C) is shown. (D) Co-incubation of isolated mitochondria from *S. cerevisiae* with the isolated Ubp3 variants with (+) or without (-) purified Bre5. Deubiquitination was assessed by SDS–PAGE and western blotting using an anti-Ubiquitin antibody. Mean ± SD of at least three independent experiments normalized to control is shown. Significance levels p< 0.05 (∗), p<0.01 (∗∗), p<0.001 (∗∗∗) or p<0.0001 (∗∗∗∗) using a two-way ANOVA test are indicated.

## Discussion

Our study shows that Ubp3 is a central deubiquitinase acting at the crossroad between ubiquitin-dependent mitophagy and remodelling of the mitochondrial ubiquitin pool. One very important finding is that mitochondria are heavily ubiquitinated under steady-state conditions and that this pool of ubiquitin is grossly depleted upon induction of autophagy and mitophagy. We could show that Ubp3, a known negative regulator of mitophagy (Müller et al., 2015), is required for this dynamic removal of ubiquitin from mitochondria *in vivo* as well as *in vitro*. This requires the catalytic activity of Ubp3 but does not require Bre5. The latter is fully in line with our recent study characterizing Ubp3 *in vitro* using various ubiquitinated substrates (Bostelmann-Arp et al., 2026). We further showed that the catalytic activity of Ubp3 is needed for inhibiting mitophagy, a finding that would be expected but was formally not shown so far. Interestingly, global mitochondrial ubiquitination does not depend on the polyubiquitin-encoding *UBI4* gene, albeit this factor is needed for mitophagy as demonstrated recently (Altin et al., 2025). Here we also showed that Ubi4-GFP is partially located to mitochondria under steady-state condition and that Ubi4 acts epistatic to Ubp3 in mitophagy. These findings together thus strongly suggest that Ubi4 is required for ubiquitination of specific substrates at the mitochondrial surface to mediate mitophagy, rather than to be essential for the observed extended mitochondrial ubiquitination. Moreover, Ubi4 is also not needed for mitochondrial deubiquitination as opposed to Ubp3.

All this raises the important question: What could be the physiological role of Ubp3-dependent mitochondrial deubiquitination under conditions promoting mitophagy? Based on the reported inhibitory role in mitophagy and on the known dynamic translocation of the Ubp3/Bre5 complex to mitochondria during mitophagy, we have earlier speculated that Ubp3 limits excessive mitophagy and thus ensures that not all mitochondria are degraded under stress (Müller et al 2015; Behrendt et al.). Thereby, Ubp3 is a critical factor ensuring mitochondrial maintenance as this endosymbiont-derived organelle cannot be formed *de novo*. Mechanistically, Ubp3 would at least deubiquitinate specific mitochondrial outer membrane proteins required for mitophagy. This view is fully in-line with our recent finding that ubiquitination of Fzo1, an outer membrane protein needed for mitochondrial fusion, is required for mitophagy and that Fzo1 is degraded during mitophagy in a proteasome- and *UBI4*-dependent manner (Altin et al., 2025). Thus, at least one factor needed for ubiquitin-dependent mitophagy, namely Fzo1, is apparently ubiquitinated at the mitochondrial surface. Since global deubiquitination of mitochondria is apparently not sufficient to limit mitophagy, it is likely that a restricted subset of very specific ubiquitinated receptors underlie the reported mechanism of CxUb/UbN77-mediated mitophagy (Altin et al., 2025). In this scenario, Ubp3 acts by deubiquitinating not only globally at the mitochondrial surface but specifically at a subset of receptors, thereby limiting mitophagy. The epistatic relationship between Ubi4 and Ubp3 places the function of Ubi4 upstream of Ubp3 in mitophagy regulation. It is thus important to separate the dynamic global mitochondrial ubiquitination profile from specific substrates involved in mitophagy. Identifying mitochondrial substrates whose (de-)ubiquitination is specifically regulated by Ubi4 and Ubp3 will thus be a crucial step for understanding the role of ubiquitin dynamics in mitophagy.

Thus, our observation that Ubp3 mediates deubiquitination of basically the entire pool of ubiquitinated mitochondrial proteins raises another additional possible role of Ubp3 beyond regulating mitophagy. We propose that Ubp3-dependent global deubiquitination of mitochondria also ensures the release of a ubiquitin pool that can promote other ubiquitin-dependent cellular quality control pathways, including the ubiquitin-proteasome-system (UPS) and various forms of autophagy. Given that mitochondria occupy roughly 20 to 30% of the cell volume, this pool would likely represent a substantial source of ubiquitin. Future studies will need to dissect this further.

Notably, Ubp3 and Ubp4/Doa4 were the only prominent DUBs apparently affecting mitochondrial deubiquitination after inhibition of TORC1 signalling identified in our screen (Fig. 2A). Since *S. cerevisiae* contain only about 20 of DUBs with a constrained array of specialized functions, it is reasonable that there are enzymes that have evolved a strong pleiotropy upon specific types of stress. *Δubp4* cells accumulated high amounts of ubiquitinated mitochondrial proteins both under control conditions (data not shown) as well as after treatment with rapamycin (Fig. 2). However, cells lacking Ubp4/Doa4 had higher overall ubiquitination of mitochondrial proteins but still showed deubiquitination upon treatment with rapamycin (data not shown) suggesting that Ubp4 is not required directly for deubiquitinating mitochondria like Ubp3. This observation was still unexpected as Ubp4 was earlier reported to cause a general depletion of the ubiquitin pool which was attributed to impaired ubiquitin recycling in the UPS (Swaminathan et al., 1999). Two effects of Ubp4/Doa4 deletion could possibly explain this shift in general ubiquitin dynamics. First, *UBI4* mRNA level were reported to be upregulated in cells lacking Ubp4 (Swaminathan et al., 1999) and could simply provide increased ubiquitin levels that are partially also stored at mitochondria, resulting in increased mitochondrial ubiquitination. Second, lack of Ubp4/Doa4 was reported to result in accumulation of substrates that cannot be further processed (Papa et al., 1999; Swaminathan et al., 1999), which could also account for ubiquitinated mitochondrial cargo. Nevertheless, future studies will have to clarify why there is a shift toward mitochondrial ubiquitination in cells lacking Ubp4. Also, *Δubp6* cells showed a severe phenotype in which mitochondria were significantly less ubiquitinated treated with or without rapamycin (Fig. 2B). Since Ubp6 is a proteasome-associated DUB essential for recycling of ubiquitin molecules (Hanna et al., 2006; Hung et al., 2022), loss of Ubp6 can lead to a general ubiquitin depletion and thus decreased ubiquitination capability also at mitochondria.

The role of Bre5 in remodelling mitochondrial ubiquitination remains was also addressed here. Our findings show that Bre5 is neither needed for a global deubiquitination *in vivo* (Fig. 2C), nor for deubiquitination of mitochondria *in vitro* (Fig. 4). Our findings here and reported earlier (Bostelmann-Arp et al., 2026) show that Bre5 is not strictly needed as a cofactor for the deubiquitination efficacy of Ubp3, but rather might stabilizes Ubp3 *in vivo* and serves in defining substrate specificity, including the receptors targeted by Ubi4-mediated ubiquitination thereby promoting the role of Ubp3 in mitophagy inhibition. This would be in line with earlier research from our group and connect the findings showing a role of Bre5 in Ubp3-directed inhibition of mitophagy (Müller et al., 2015) with our recent discovery that Ubp3 can deubiquitinate proteins without the presence or binding of its proposed cofactor Bre5 (Bostelmann-Arp et al., 2026).

Mitochondrial quality control and the ubiquitin system are associated with the development of diseases in humans, including cancer and neurological disorder such as Parkinson’s disease. Whether an orthologous mitochondria-associated DUB in humans plays a similar role as Ubp3 remains to be investigated. The human ortholog of Ubp3, USP10, represents an interesting candidate considering its established links to cellular energy metabolism (Deng et al., 2016). However, the landscape of mammalian DUBs is considerably more complex, making these future analyses much more complex.

Collectively, our data support a model in which Ubp3 contributes to global mitochondrial ubiquitin dynamics and remodels ubiquitination upon induction of mitophagy. Ubp3-mediated editing of ubiquitinated mitochondrial proteins is involved in, but not limited to, blocking Ubi4-dependent mitophagy. Instead, our findings suggest that a restricted subset of Ubi4-dependent receptors or very specific ubiquitin linkages are likely underlying the mechanism of mitophagy regulation while global ubiquitin availability is unlikely to account primarily for mitophagy regulation. Here, we put forward the hypothesis that the vast amount of ubiquitinated mitochondrial proteins act as a storage platform and Ubp3 is involved in supplying the cell with free ubiquitin molecules upon stress. Defining the exact substrates involved and determining how Ubp3 and possibly Bre5 are recruited to these mitochondrial targets will be crucial for understanding the connection between ubiquitin dynamics and cellular quality control.

## Methods

### Cell cultivation and general experimental procedures

All *S. cerevisiae* strains used in this study are listed in Table S1. Single knockout strains were obtained from the EUROSCARF library, with a MATα background (BY4742: his3Δ1; leu2Δ0; lys2Δ0; ura3Δ0) in which the respective gene was replaced with a KanMX module. The Ubi4-GFP fusion strain was generated by homologous recombination using a GFP-coding plasmid as previously described (Gardner and Jaspersen, 2014) in the BY4741 Mata background (his3Δ1; leu2Δ0; met15Δ0; ura3Δ0). To stain mitochondria plasmid-mediated mtDsRED expression was used. Table S2 lists the primers and plasmids used for gene manipulation. Transformation was done using the chemical transformation protocol in TE buffer containing lithium acetate/PEG and recovery followed in selection medium containing either 200 µg/ml G418 or lacking amino acids (-HIS) and/or leucin (-LEU).

Cells were cultured using standard methods at growth temperature of 30 °C. Incubation occurred in YP (1 % yeast extract, 2 % peptone) medium with 2 % glucose (YPD) or 2 % glycerol (G), or in synthetic minimal medium with 2 % glucose lacking the amino acid histidine (-HIS) and/or leucin (-LEU) for selection of auxotrophic mutants. For induction of mitophagy, strains were grown in YPG medium and treated with 1 µM rapamycin (+) for the indicated time, or left untreated (-).

### Microscopy

The BY4741 Mata Ubi4-GFP-HIS3MX6 cells expressing mitochondrial RFP (mitoDsRed) were cultured in YPG medium at 30 °C and treated for 0, 1, 5 or 24 hours with 1 µM rapamycin before gentle collection of 8 OD by centrifugation at 1000 g for 1 min at room temperature. Cell pellet was resuspended in 1 ml YPG medium. Subsequently, 2 µl cell suspension were mixed with 2 µl of low melting agarose (2 % in dH_2_O) and transferred onto a microscopy slide. Images were acquired using a Zeiss Axio Observer.D1 fluorescence microscope with a 63×/1.44 Plan-Apochromat immersion oil objective and accompanying ZEN 2012 software.

### Subcellular fractionation

Designated *S. cerevisiae* strains were grown in YPG medium and expanded to 1 litre of volume by ongoing dilutions of the cultures over three days to maintain them in a non-stationary growth phase. Cultures were then treated with 1 µM of rapamycin for the indicated time or left untreated as control. Cells were collected by centrifugation, treated with DTT and converted to protoblasts using Zymolyase (Amsbio). The cells were gently homogenized using a Douncer followed by differential centrifugation to obtain cytosolic, nuclear and crude mitochondrial fractions. For the ubiquitin dynamics shown in Figure 1 and the DUB screen in Figure 2, crude mitochondria were used. In all other cases crude mitochondria were further purified using sucrose gradient density centrifugation. All fractions including 200 µl of the total lysate were either immediately used for downstream analysis or snap-frozen and stored at -80 °C. *S. pombe* cells were cultivated as described in (Moreno et al., 1991). *U. maydis* cells were cultured as described before (Postma et al., 2026) and protoblasts were obtained using Vinotaste PRO (Novozyme, Denmark).

### SDS PAGE western blotting

Protein concentration of samples was determined using the BCA assay from ThermoScientific (ref) and equal amounts of 20 µg protein per lane were diluted with dH_2_O to 15 µl to which 5 µl 4X Laemmli buffer was added. For western blotting analysis, proteins were resolved by Tris/Glycine (12%) SDS-PAGE, and transferred onto nitrocellulose membranes. For protein detection, several antibodies were used, in 5% milk in TBST, the buffer also used during the washing steps. Bmh2 was detected with a rabbit polyclonal antibody, kindly gifted by Walter Neupert (LMU Munich, Germany), diluted to 1:1000. The rabbit polyclonal antibody against *S. cerevisiae* Sec61 (used at 1:1000) was a kind gift from Richard Zimmermann. To detect mtALP, we used a polyclonal antibody raised in rabbit against the fusion protein MBP-Pho8, and to detect mitochondrial protein Tim44, a polyclonal antibody raised in rabbit was used, both diluted to 1:1000. The following commercial antibodies were used: anti-GFP (Sigma, from mouse IgG, clone 7.1, used at 1:1000), anti-Porin (Abcam (ab14734), from mouse IgG, clone 20B12AF2, used at 1:1000), and anti-Ubiquitin (Sigma, monoclonal antibody from mouse IgG, clone P4D1-A11, used at 1:1000).

### Mitophagy assay

The alkaline phosphatase (ALP) assay was used to quantify mitophagy in yeast strains expressing the mtALP reporter, as described previously (Mendl et al., 2011). Briefly, ALP activity was determined by measuring the absorption of p-nitrophenyl phosphate at 405 nm after 15 min, using a Tecan plate reader. Protein concentrations were determined by carrying out the BCA assay according to standard protocol (ThermoScientific). Specific activities were normalized to the activity of the indicated control strain. All strains were deleted for the endogenous alkaline phosphatase PHO8. In addition, cells were harvested and cell lysates analysed by SDS-PAGE and western blotting as described above.

### Protein expression and purification

The full-length proteins Bre5 and Ubp3 (FL), as well as the catalytic domain of Ubp3 (CD) without or with the catalytically inactivating mutation (CD) were expressed and purified as described by earlier (Bostelmann-Arp et al., 2026). Briefly, the genes from *S. cerevisiae* and their truncation and mutant variants were cloned into a pQlink plasmid by Gibson assembly. The resulting constructs were expressed in *Escherichia coli (E. coli)* and overexpression was induced using IPTG. For purification, cell pellets were harvested and lysed, followed by affinity chromatography. Proteins were further purified by size exclusion chromatography (SEC) and concentrated. The purified proteins were either immediately used for downstream experiments or snap-frozen and stored at -80 °C.

### DUB assay

For the deubiquitination assay of isolated mitochondria ex vivo shown in Figure 4A, 20 µg of mitochondria were mixed with varying concentrations of 0, 50, 100, 150, 300 or 600 nM purified full-length Ubp3 in a total volume of 15 µl and incubated for 1 hour at 30 °C. For the other assays, purified proteins Bre5 and the Ubp3 variants (FL, CD, and CD) were incubated at 150 nM. The reaction was stopped by immediately adding 5 µl 4X Laemmli and boiling the probes for 5 minutes at 95 °C followed by western blotting to detect changes in total ubiquitination.

### Data analysis

Quantifications of fluorescence microscopy images, mtALP activity and proteins levels was performed using Fiji and GraphPad Prism 8. Protein quantities were normalized to Ponceau staining and the indicated loading controls. Statistical details can be found in the corresponding figure legend. Significance was calculated using two-way ANOVA and p-values are displayed as ns with p>0.05, * with p<0.05, ** with p<0.01, *** with p<0.001 and **** with p<0.0001.

## Supporting information

SFigure 1

Supplementary Material STab1 STab2 SFig1Legend

## Acknowledgements

We would like to thank Sandra Schmidt for technical support. We thank Kathleen Gould for the *S. pombe* strain and Claudine Kraft for Ubp3-plasmids. This work is part of the collaboration research center SFB1535 (projects A04, A03, and A02 to ASR, LS, MF, and UF), funded by the Deutsche Forschungsgemeinschaft (DFG, German Research Foundation)-SFB 1535 Project ID 458090666.

## References

Abeliovich, H. 2023. Mitophagy in yeast: known unknowns and unknown unknowns. Biochem J. 480:1639–1657.

Altin, S., T. Simoes, C. Behrendt, V. Anton, D. Domke, K.M. Völtzke, R. Kumar, H. Nolte, T. Hermanns, N. Brocke-Ahmadinejad, K. Bendrin, M. Zimmermann, R. Buttner, N. Ventura, M. Kruger, R.J. Dohmen, K. Hofmann, T. Hoppe, A.S. Reichert, and M. Escobar-Henriques. 2025. Ubiquitin precursor with C-terminal extension promotes proteostasis and longevity. Molecular cell. 85:3677–3693 e3677.

Anton, F., G. Dittmar, T. Langer, and M. Escobar-Henriques. 2013. Two deubiquitylases act on mitofusin and regulate mitochondrial fusion along independent pathways. Molecular cell. 49:487–498.

Behrendt, C., and A.S. Reichert. 2016. Mitophagy and deubiquitination in yeast - the power of synthetic quantitative array technology. Molecular & cellular oncology. 3:e1038422.

Biswas, S., and P. D’Silva. 2025. Ubp2 modulates DJ-1-mediated redox-dependent mitochondrial dynamics in Saccharomyces cerevisiae. PLoS genetics. 21:e1011353.

Bostelmann-Arp, L., S. Khosa, J. Reiners, K. Mayor Völtzke, S.H.J. Smits, A.S. Reichert, and L. Schmitt. 2026. In vitro characterization of the baker’s yeast deubiquitinase Ubp3. bioRxiv:2026.2008.2019.745719.

Deng, M., X. Yang, B. Qin, T. Liu, H. Zhang, W. Guo, S.B. Lee, J.J. Kim, J. Yuan, H. Pei, L. Wang, and Z. Lou. 2016. Deubiquitination and Activation of AMPK by USP10. Molecular cell. 61:614–624.

Dikic, I. 2017. Proteasomal and Autophagic Degradation Systems. Annual review of biochemistry. 86:193–224.

Gardner, J.M., and S.L. Jaspersen. 2014. Manipulating the yeast genome: deletion, mutation, and tagging by PCR. Methods in molecular biology. 1205:45–78.

Hanna, J., N.A. Hathaway, Y. Tone, B. Crosas, S. Elsasser, D.S. Kirkpatrick, D.S. Leggett, S.P. Gygi, R.W. King, and D. Finley. 2006. Deubiquitinating enzyme Ubp6 functions noncatalytically to delay proteasomal degradation. Cell. 127:99–111.

Hung, K.Y.S., S. Klumpe, M.R. Eisele, S. Elsasser, G. Tian, S. Sun, J.A. Moroco, T.C. Cheng, T. Joshi, T. Seibel, D. Van Dalen, X.H. Feng, Y. Lu, H. Ovaa, J.R. Engen, B.H. Lee, T. Rudack, E. Sakata, and D. Finley. 2022. Allosteric control of Ubp6 and the proteasome via a bidirectional switch. Nature communications. 13:838.

Iesmantavicius, V., B.T. Weinert, and C. Choudhary. 2014. Convergence of ubiquitylation and phosphorylation signaling in rapamycin-treated yeast cells. Molecular & cellular proteomics : MCP. 13:1979–1992.

Kanki, T., D.J. Klionsky, and K. Okamoto. 2011. Mitochondria autophagy in yeast. Antioxid Redox Signal. 14:1989–2001.

Kinner, A., and R. Kölling. 2003. The yeast deubiquitinating enzyme Ubp16 is anchored to the outer mitochondrial membrane. FEBS letters. 549:135–140.

Komander, D., and M. Rape. 2012. The ubiquitin code. Annual review of biochemistry. 81:203–229.

Kondadi, A.K., and A.S. Reichert. 2024. Mitochondrial Dynamics at Different Levels: From Cristae Dynamics to Interorganellar Cross Talk. Annu Rev Biophys. 53:147–168.

Leo, M., G. Fanelli, S. Di Vito, B. Traversetti, M. La Greca, R.A. Palladino, A. Montanari, S. Francisci, and P. Filetici. 2018. Ubiquitin protease Ubp8 is necessary for *S. cerevisiae* respiration. Biochim Biophys Acta Mol Cell Res.

Mendl, N., A. Occhipinti, M. Muller, P. Wild, I. Dikic, and A.S. Reichert. 2011. Mitophagy in yeast is independent of mitochondrial fission and requires the stress response gene WHI2. Journal of cell science. 124:1339–1350.

Moreno, S., A. Klar, and P. Nurse. 1991. Molecular genetic analysis of fission yeast Schizosaccharomyces pombe. Methods in enzymology. 194:795–823.

Müller, M., P. Kötter, C. Behrendt, E. Walter, C.Q. Scheckhuber, K.D. Entian, and A.S. Reichert. 2015. Synthetic quantitative array technology identifies the Ubp3-Bre5 deubiquitinase complex as a negative regulator of mitophagy. Cell reports. 10:1215–1225.

Narendra, D.P., S.M. Jin, A. Tanaka, D.F. Suen, C.A. Gautier, J. Shen, M.R. Cookson, and R.J. Youle. 2010. PINK1 is selectively stabilized on impaired mitochondria to activate Parkin. PLoS Biol. 8:e1000298.

Papa, F.R., A.Y. Amerik, and M. Hochstrasser. 1999. Interaction of the Doa4 deubiquitinating enzyme with the yeast 26S proteasome. Mol Biol Cell. 10:741–756.

Pereira, C., V. Costa, L.M. Martins, and L. Saraiva. 2015. A yeast model of the Parkinsons disease-associated protein Parkin. Exp Cell Res.

Postma, J., P. Künzel, N. Wierckx, K. Schipper, and M. Feldbrügge. 2026. Synchronous Induction and Microscopic Analysis of Hyphal Growth Using Laboratory Strain AB33. In Ustilago maydis: Methods and Protocols. A.N.T. Phan, L. Blank, and J. Schirawski, editors. Springer US, New York, NY. 83–94.

Sheng, X., Z. Xia, H. Yang, and R. Hu. 2024. The ubiquitin codes in cellular stress responses. Protein & Cell. 15:157–190.

Swaminathan, S., A.Y. Amerik, and M. Hochstrasser. 1999. The Doa4 deubiquitinating enzyme is required for ubiquitin homeostasis in yeast. Mol Biol Cell. 10:2583–2594.

Vives-Bauza, C., C. Zhou, Y. Huang, M. Cui, R.L. de Vries, J. Kim, J. May, M.A. Tocilescu, W. Liu, H.S. Ko, J. Magrane, D.J. Moore, V.L. Dawson, R. Grailhe, T.M. Dawson, C. Li, K. Tieu, and S. Przedborski. 2010. PINK1-dependent recruitment of Parkin to mitochondria in mitophagy. Proceedings of the National Academy of Sciences of the United States of America. 107:378–383.

Youle, R.J., and D.P. Narendra. 2011. Mechanisms of mitophagy. Nature reviews. Molecular cell biology. 12:9–14.

Zimmermann, M., and A.S. Reichert. 2017. How to get rid of mitochondria: crosstalk and regulation of multiple mitophagy pathways. Biological chemistry. 399:29–45.

