## Supplementary figures and images for "Ubp3 mediates dynamic deubiquitination of mitochondria upon induction of mitophagy"

### SFigure 1

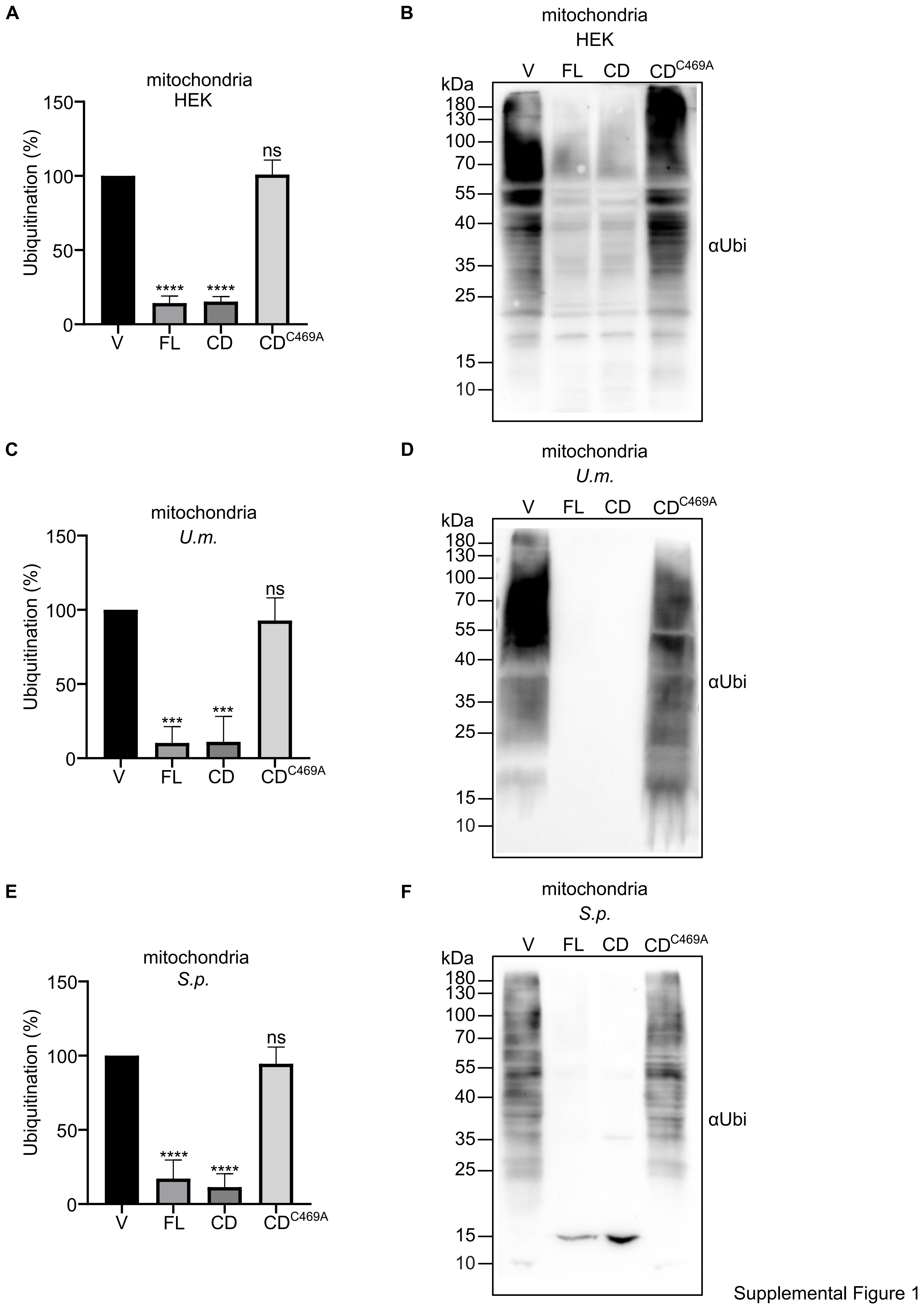


Supplemental Figure 1
