## Supplementary Material STab1 STab2 SFig1Legend for "Ubp3 mediates dynamic deubiquitination of mitochondria upon induction of mitophagy"

**Supplementary Tables and Figure legends**

**Supplementary Tables**

**Table S1. Yeast strains used in this study**

| Genotype | Source |
|---|---|
| ***S. cerevisiae* strains** | |
| BY4741 Mata ubi4-GFP-HIS3MX6 [pHS12mitoDsRed] | Bevis et al 2002 |
| BY4742 Matα his3Δ1 ; leu2Δ0 ; lys2Δ0 ; Ura3Δ0 | This study |
| BY4742 Matα ubi4::kanMX | Euroscarf |
| BY4742 Matα bre5::kanMX | Euroscarf |
| BY4742 Matα ubp1::kanMX | Euroscarf |
| BY4742 Matα ubp2::kanMX | Euroscarf |
| BY4742 Matα ubp3::kanMX | Euroscarf |
| BY4742 Matα ubp4::kanMX | Euroscarf |
| BY4742 Matα ubp5::kanMX | Euroscarf |
| BY4742 Matα ubp6::kanMX | Euroscarf |
| BY4742 Matα ubp7::kanMX | Euroscarf |
| BY4742 Matα ubp8::kanMX | Euroscarf |
| BY4742 Matα ubp9::kanMX | Euroscarf |
| BY4742 Matα ubp11::kanMX | Euroscarf |
| BY4742 Matα ubp12::kanMX | Euroscarf |
| BY4742 Matα ubp13::kanMX | Euroscarf |
| BY4742 Matα ubp14::kanMX | Euroscarf |
| BY4742 Matα ubp15::kanMX | Euroscarf |
| BY4742 Matα ubp16::kanMX | Euroscarf |
| BY4742 Matα yuh1::kanMX | Euroscarf |
| BY4742 Matα pho8::his3 [pYX242 mtPho8] | This study |
| BY4742 Matα ubi4::kanMX pho8::his3 [pYX242 mtPho8] | This study |
| BY4742 Matα bre5::kanMX pho8::his3 [pYX242 mtPho8] | This study |
| BY4742 Matα ubp3::kanMX pho8::his3 [pYX242 mtPho8] | This study |
| BY4742 Matα ubi4::kanMX pho8::his3 ubp3::natMX [pYX242 mtPho8] | This study |
| BY4742 Matα ubp3::kanMX pho8::natMX [pRS315 leer] [pVT100UmtPho8] | This study |
| BY4742 Matα ubp3::kanMX pho8::natMX [pRS315_endUBP3] [pVT100UmtPho8] | This study |
| BY4742 Matα ubp3::kanMX pho8::natMX [pRS315_endUBP3-C469A][pVT100UmtPho8] | This study |
| ***Ustilago maydis* strain** | |
| AB33 Atp3-Gfp (*a2* Pnar1::bW2, bE1/Ptef:atp3-l1-gfp-swatp3_3'UTR *atp3* (*upp3*Δ; natR)) | M. Feldbrügge |

***S. pombe* strain**

UFY605 (his3-D1, ade6-M210, leu1-32, ura4-D18, h-) K. Gould (VU, USA)

**Table S2. Plasmids used in this study**

| Name | Marker | Source |
|---|---|---|
| pYX242 mtALP | LEU2 | Mendl et al., 2011 |
| pHS12 mtDsRed | LEU2 | Mendl et al., 2011 |
| pVT100U mtALP | URA3 | Mendl et al., 2011 |
| pRS315 empty | LEU2 | Kraft et al., 2008 |
| pRS315 endosomal-Ubp3 | LEU2 | Kraft et al., 2008 |
| pRS315 endosomal-Ubp3-C469A | LEU2 | Cebollero et al., 2012 |
| pFA6-kanMX4 | G418 | Addgene 39296 |

### Supplementary Figure legends

#### Supplemental Figure 1. Ubp3 acts as a conserved remodeller of mitochondrial ubiquitination independent of the origin of mitochondria

Mitochondria isolated from human HEK cells (A and B)*, U. maydis* (C and D) or *S. pombe* (E and F) were incubated with either vehicle control (V), or purified proteins full length Ubp3 (FL), the catalytic domain of Ubp3 alone (CD), the catalytic domain of Ubp3 carrying the inactivating C469A mutation ($CD^{C469A}$). Quantifications of the blots are shown on the left (A, C and E) and a representative experiment on the right (B, D and F), respectively. Mean ± SD of at least three independent experiments normalized to control is shown. Significance levels $p < 0.05$ (∗), $p<0.01$ (∗∗), $p<0.001$ (∗∗∗) or $p<0.0001$ (∗∗∗∗) using a two-way ANOVA test are indicated.
